# Regression of Human Breast Carcinoma in Nude Mice After Ad*sflt* Gene Therapy is Mediated by Tumor Vascular Endothelial Cell Apoptosis

**DOI:** 10.1101/2020.07.19.210856

**Authors:** Angelina Felici, Donald P. Bottaro, Antonella Mangoni, Petra Reusch, Dieter Marmé, Imre Kovesdi, Dinuka M. De Silva, Young H. Lee, Maurizio C. Capogrossi, Judith Mühlhauser

## Abstract

Two vascular endothelial growth factor (VEGF) receptors, FLT-1 and KDR, are expressed preferentially in proliferating endothelium. There is increasing evidence that recombinant, soluble VEGF receptor domains interfering with VEGF signaling may inhibit *in vivo* neoangiogenesis, tumor growth and metastatic spread. We hypothesized that a soluble form of FLT-1 receptor (sFLT-1) could inhibit the growth of pre-established tumors via an anti-angiogenic mechanism. A replication-deficient adenovirus (Ad) vector carrying the sflt-1 cDNA (Ad*sflt*) was used to overexpress the sFLT-1 receptor in a breast cancer animal model. MCF-7 cells, which produce VEGF, were used to establish solid tumors in the mammary fat pads of female nude mice. After six weeks tumors were injected either with Ad*sflt*, or a negative control virus (AdCMV.βgal). After six months, average tumor volume in the Ad*sflt-*infected group (33 ± 22 mm3) was decreased by 91% relative to that of the negative control group (388 ± 94 mm3; *P*<0.05). Moreover, 10 of 15 Ad*sflt-*infected tumors exhibited complete regression. The vascular density in Ad*sflt-*infected tumors was reduced by 50% relative to that of negative controls (*P*<0.05), consistent with sFLT-1-mediated tumor regression through an anti-angiogenic mechanism. Moreover, cell necrosis and fibrosis associated with long-term regression of Ad*sflt*–infected tumors were preceded by apoptosis of tumor vascular endothelial cells. Mice treated with Ad*sflt* intratumorally showed no delay in the healing of cutaneous wounds, providing preliminary evidence that Ad-mediated sFLT-1 overexpression may be an effective anti-angiogenic therapy for cancer without the risk of systemic anti-angiogenic effects.

## Introduction

Angiogenesis is required for solid tumor growth, and inhibition of angiogenesis has been proposed as a potential strategy for the treatment of cancer (Folkman J, 1995; Harris *et al*., 1996). Among the many paracrine angiogenic growth factors produced by cancer cells, vascular endothelial growth factor (VEGF) appears to play a key role in the growth of a variety of tumors (Senger *et al*., 1993; Shibuya M, 1995). A 46-kDa secreted polypeptide, VEGF binds the fms-like tyrosine kinase receptor FLK-1, the mouse homolog of KDR (Terman *et al*., 1992; Millauer *et al*., 1993), as well as FLT-1 (Shibuya *et al*., 1990; De Vries *et al*., 1992), which are expressed almost exclusively on endothelial cells. The importance of VEGF receptors in angiogenesis is underscored by gene deletion and transgenic animal studies where disruption of the VEGF/FLK-1 and VEGF/FLT-1 pathways leads to early embryonic lethality due to abnormalities in the embryonic vasculature (Fong *et al*., 1995; Shalaby *et al*., 1995).

Several inhibitors of angiogenesis with normal physiologic roles have been identified. Among these are thrombospondin-1 which modulates endothelial cell adhesion, motility and proteolytic activity by sequestering angiogenesis inducers (Dameron *et al*., 1994), and platelet factor 4 which inhibits endothelial cell growth *in vitro* and tumor associated angiogenesis *in vivo* (Tanaka *et al*., 1997). The proteolytic protein fragments angiostatin (O’Reilly *et al*., 1994) and endostatin (O’Reilly *et al*., 1997) inhibit endothelial cell proliferation, angiogenesis and tumor growth. Furthermore, tissue inhibitors of metalloproteinases, TIMP-1 and TIMP-2, inhibit collagenase activity affecting endothelial cell migration and proliferation (Moses and Langer, 1991; Ray and Stetler-Stevenson, 1994). Thus, normal angiogenesis appears to be tightly regulated through the balance of inhibitors and inducers, and increased inducer activity can lead to tumor vascularization and tumor growth (Hanahan and Folkman, 1996).

Many early studies showed the potential therapeutic utility of pharmacological angiogenesis inhibitors, such as fumagillin or thalidomide (Ingber *et al*., 1990; D’Amato *et al*., 1994), and those targeting the VEGF signaling pathway to combat pathological angiogenesis. Inhibition of VEGF action by specific anti-VEGF antibodies can block ligand binding, angiogenesis and tumor growth in immundeficient mice (Kim *et al*., 1993; Shih and Lindley, 2006). A VEGF variant has been shown to antagonize VEGF-stimulated receptor phosphorylation and endothelial cell proliferation (Siemeister *et al*., 1998), and a dominant negative FLK-1 receptor mutant inhibited tumor growth *in vivo* (Millauer *et al*., 1994). Recombinant soluble VEGF receptor domains interfering with VEGF signaling exert inhibitory effects on *in vivo* neoangiogenesis (Aiello *et al*., 1995), tumor growth (Lin *et al*., 1998; Sallinen *et al*., 2009) and metastatic spread (Kong *et al*., 1998). Among these antagonists, the soluble form of the FLT-1 receptor (sFLT-1) shows promise as a potent inhibitor of endothelial cell function (Kendall and Thomas, 1993) and angiogenesis (Aiello *et al*., 1995; Kong *et al*., 1998; Bazan-Peregrino *et al*., 2013). This truncated FLT-1 isoform is produced by alternative splicing of flt-1 mRNA (Kendall and Thomas, 1993), and is comprised of the first six extracellular immunoglobin (Ig)-like domains of the full-length receptor and a specific 31 amino acid carboxy terminus.

Virotherapy provides a novel approach to cancer treatment. Oncolytic viruses have been shown to infect and damage cancerous cells and are generally unable to replicate in normal cells. While variants of the VEGF ligand and receptor proteins have much to offer as reagents for anti-angiogenesis therapy, their overall efficacy is partly contingent on solutions to pharmacological problems common to biologicals, such as delivery and stability. In this regard, gene therapy using recombinant adenoviral vectors represents a promising alternative approach; such vectors have been shown to efficiently deliver and express genes in a wide variety of tissues including solid tumors (for review see Descamps *et al*., 1996; Kong and Crystal, 1998). Moreover, their local application is less toxic than pharmacological substances which often exhibit strong and/or undesired side effects. Additional advantages to these and other viral and non viral systems are the high efficiency of cell infection, and the limited spatial and temporal expression of the transgene. This feature may be particularly important for anti-angiogenic therapy, to avoid inhibition of angiogenic activities that are part of normal homeostasis.

The present study was carried out to investigate the therapeutic potential and mechanism of action of adenovirus-mediated sflt-1 gene transfer in an MCF-7 breast carcinoma animal model. A recombinant adenovirus vector carrying the human sflt-1 cDNA was injected into pre-established MCF-7 tumors in nude mice, and tumor development was observed over a period of six months. The results demonstrate that a single injection of the sFLT-1 adenoviral vector (Ad*sflt*) significantly inhibited breast tumor growth, and led to long-term tumor regression. Our observation that sFLT-1 paracrine expression induced apoptosis of tumor vascular endothelial cells and subsequent blood vessel regression also provides new insight into the mechanism of sFLT-1 anti-angiogenic action.

## Results

### sFLT-1 Overexpression Inhibits VEGF-induced Endothelial Cell Proliferation in vitro

To test the ability of Ad*sflt* to inhibit VEGF-mediated mitogenic signaling, the proliferation of Ad*sflt*-infected HUVEC was studied in the presence of added VEGF. The results showed that human recombinant VEGF (10 ng/ml) significantly stimulated the proliferation of AdCMV.βgal-infected and uninfected HUVEC (Figure 1A). In contrast, Ad*sflt*-infected HUVEC did not respond to the addition of VEGF and exhibited a growth rate similar to that of uninfected HUVEC grown in the absence of VEGF. The Ad*sflt*-mediated inhibition of VEGF-stimulated HUVEC growth was 37.2 ± 4%, 49.9 ± 0.1% and 53.5 ± 2.9% (% inhibition of total proliferation) at 3, 4 and 5 days after Ad*sflt* infection respectively (*P<*0.05 at all time points). In contrast, bFGF-induced HUVEC proliferation was not inhibited by Ad*sflt* infection (data not shown). Overexpression of sflt-1 transgene under the experimental conditions mentioned above was assessed by Northern analysis; a striking increase of sflt-1 mRNA in Ad*sflt*-infected HUVEC was detected as early as 1 day after infection and 4 days thereafter (Figure 1B). Taken together these data show that the Ad-mediated sflt-1 gene overexpression in HUVEC leads to potent and specific inhibition of VEGF-stimulated endothelial cell growth.

**Figure 1.**
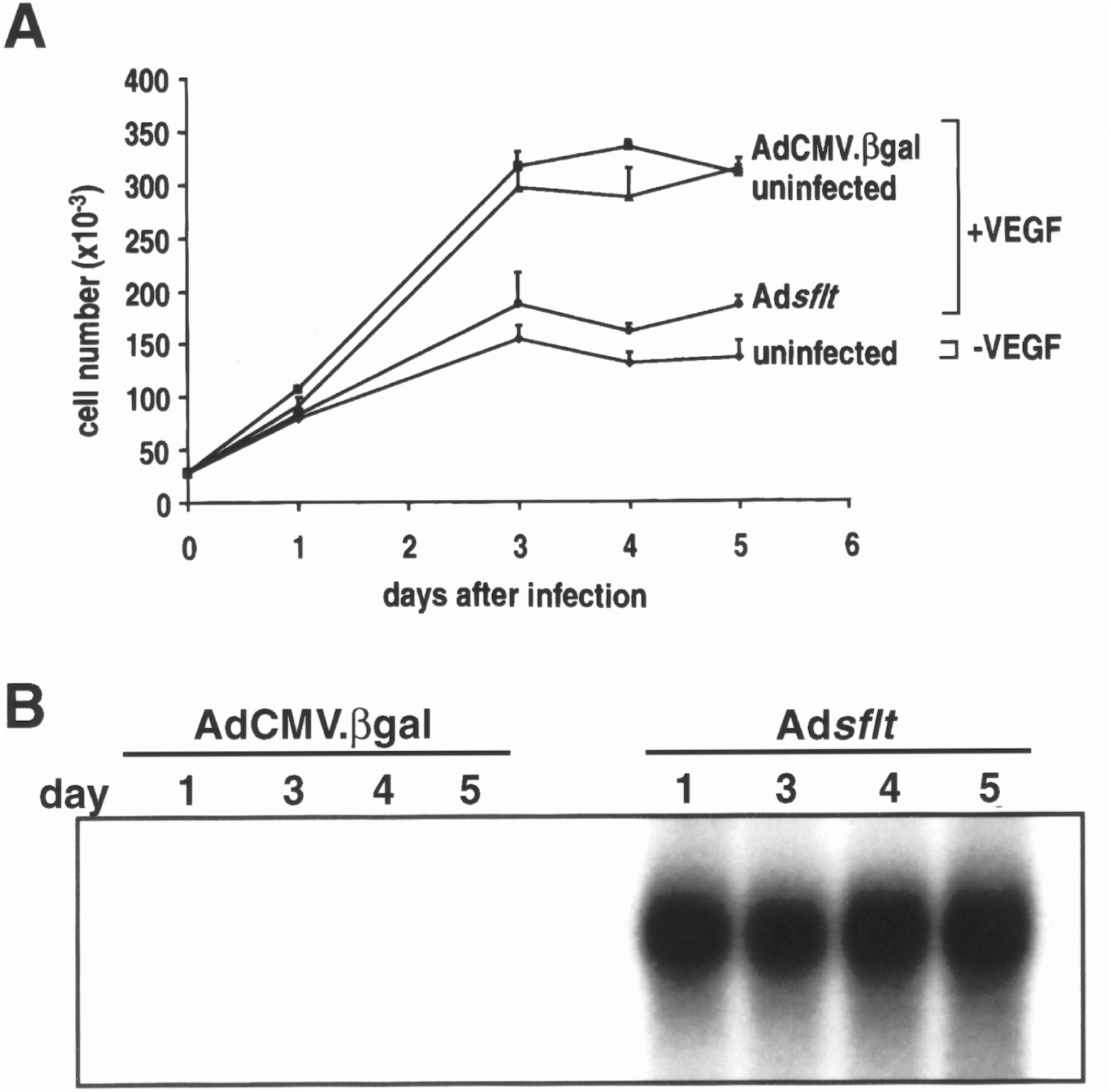
Ad*sflt*-mediated sFLT-1 overexpression inhibits VEGF-induced HUVEC proliferation *in vitro*. **(A)** Effect of Ad*sflt* infection on HUVEC proliferation *in vitro*. HUVEC (2.5×10^4^) grown in the presence of VEGF (10 ng/ml) were uninfected (triangles), or infected with 25 pfu/cell of AdCMV.βgal (squares) or Ad*sflt* (circles). Uninfected HUVEC grown in the absence of VEGF (diamonds) were used as negative control. Note the dramatic decrease in the proliferation of Ad*sflt*-infected cells grown in the presence of VEGF. The number of viable cells was determined by cell counting with a hemocytometer in triplicate. Data are presented as mean ± SE of triplicate samples of 2 experiments. **(B)** Northern analysis demonstrating that sflt-1 mRNA content was markedly increased in Ad*sflt*-*vs* AdCMV.βgal-infected HUVEC as early as 1 day after infection, and remained elevated up to day 5 after infection.

### Adsflt Inhibits Tumor Growth in vivo and Induces Long-Term Tumor Regression

VEGF is a potent tumor angiogenic factor that is produced by most tumor cells and by MCF-7 cells in culture (Hyder *et al*., 1998). Basal levels of VEGF mRNA are characteristic for established MCF-7 tumors in the nude mouse model used here (McLeskey *et al*., 1998). To determine the effect of sflt-1 gene transfer on pre-established tumor growth and angiogenesis, a single dose (2×109pfu) of Ad*sflt* vector, control Ad virus, or saline solution was administered into MCF-7 breast tumors in nude mice, and the animals were monitored for tumor growth. Briefly, well established palpable tumors (10.7 ± 1.7 mm3) were directly injected with Ad*sflt* (n=15), AdCMV.ßgal (n=14) or with saline solution (n=6), measured weekly with a caliper along the three axes (a, b and c), and their volumes were calculated (tumor volume = axbxc) (Figure 2A). The results showed modest inhibition of tumor growth in Ad*sflt*-treated tumors compared to saline-treated tumors 7 weeks post-injection (*P<*0.05), and a significant decrease in tumor size in Ad*sflt-*treated tumors *vs* saline-or AdCMV.ßgal-injected tumors (*P<*0.05) by week 9 and thereafter (Figure 2A). Fifteen weeks post-injection, 2 saline-injected tumors were removed because they were beginning to ulcerate the overlying skin. The experiment was terminated 26 weeks after treatment, because of the increasing number of ulcerating tumors, both in the saline group (2/4; 50%) and in the AdCMV ßgal-treated group (2/14; 14.2%) (Figure 2B). Between weeks 6 and 15 the control virus showed a modest but statistically significant inhibitory effect on tumor growth, possibly due to the marked immunogenicity of the βgal transgene documented previously (Lampson *et al*., 1993). However, this effect was temporary compared to the Ad*sflt* therapeutic effect. Twenty six weeks post-injection the mean volume of the tumors was significantly reduced in the Ad*sflt*-treated group (33 ± 22 mm3; n=15) compared to the saline group (323 ± 34 mm3; n=4) or the AdCMVßgal-treated group (388 ± 94 mm3; n=14). While 16 of 18 tumors (88.8%) in the control group showed steadily increased tumor size, only 2 of 15 tumors (13.3%) in the Ad*sflt*-treated group continued to enlarge 26 weeks post treatment. Moreover, at the end of the experiment 10 of 15 tumors (66.6%) injected with Ad*sflt*, compared to 2 of 18 tumors (11.1%) in the control groups, showed complete regression and their size was no longer measurable (*P<*0.05). Eight out of 10 tumors which underwent complete regression in the Ad*sflt* group had already regressed by week 9, and these animals remained tumor-free throughout the experiment.

**Figure 2.**
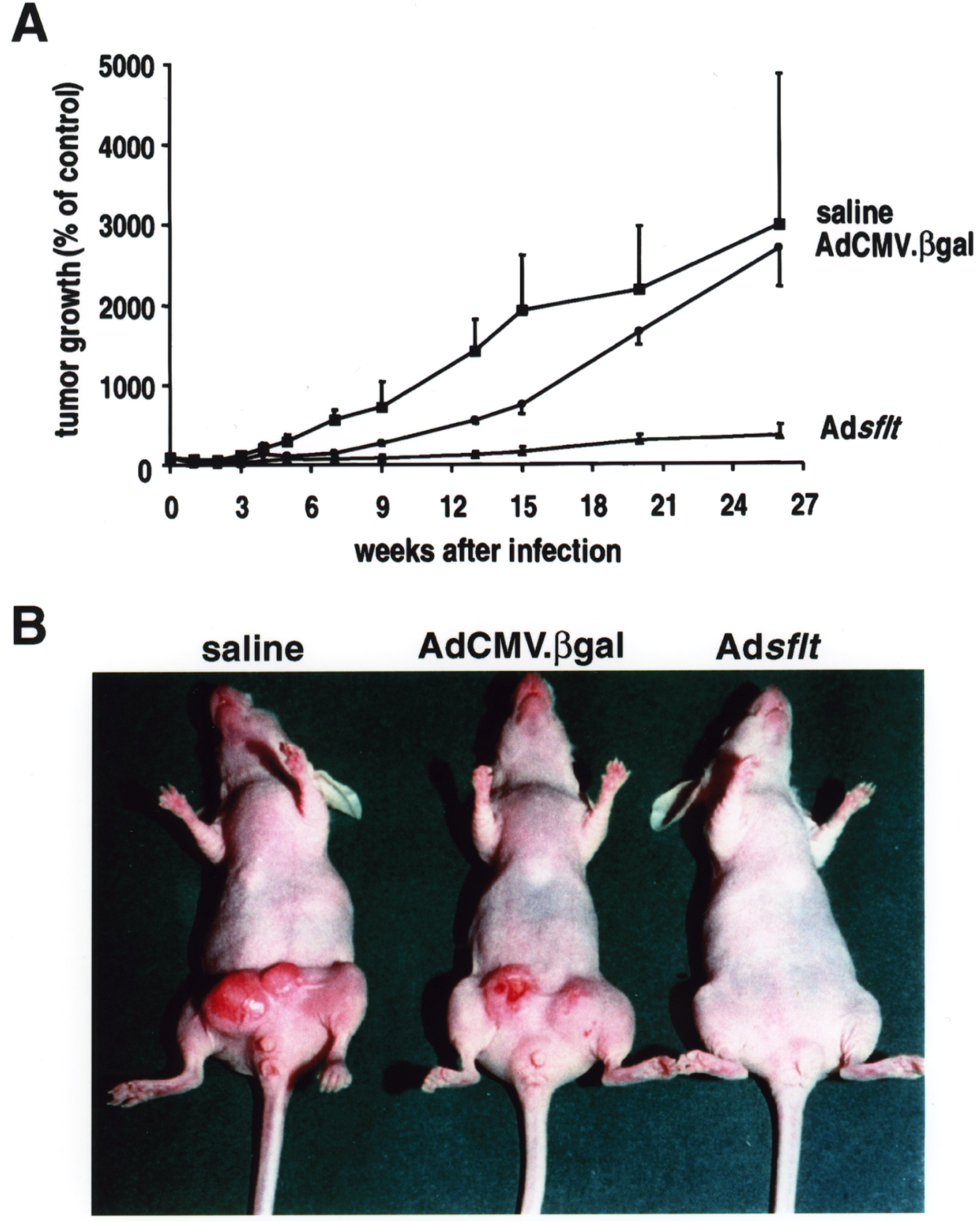
Effect of Ad*sflt* single injection on MCF-7 tumor growth in nude mice. **(A)** Inhibition of mammary tumor growth after a single administration of Ad*sflt* vector. Well-established breast tumors arising from orthotopically implanted MCF-7 cells (1×106) were injected (day 0) with Ad*sflt* (2×109 pfu/50 µl) (triangles), AdCMV.βgal (2×109 pfu/50 µl) (circles) or with saline solution (50 µl) (squares) Tumor volume was measured weekly with calipers; at each time point the volume of a single tumor was expressed as the percentage of the volume of the same tumor (100%), considered as reference at day 0. Data are presented as mean % ± SE (saline n=6; AdCMV.βgal n=14; Ad*sflt* n=15). Note that from week 9 onward there was a significant inhibition of tumor growth in Ad*sflt*-treated tumors compared to controls (*P<*0.05). **(B)** Macroscopic appearance of saline-, AdCMV.ßgal-and Ad*sflt*-treated tumors, 26 weeks after a single injection in nude mice. While control breast tumors reached a remarkable volume and were often ulcerated, Ad*sflt*-treated tumors regressed to a non-measurable size.

Histological analysis of MCF-7 tumors showed that they were well-established and vascularized before treatment (Figure 3A). Similarly, 26 weeks after AdCMV.ßgal infection, mammary tumors still exhibited a well developed peri-tumor hypervascular border zone (Figure 3B). In contrast, as shown in Figure 3C, tissue sections from tumors excised 26 weeks after treatment demonstrated that Ad*sflt*-infected tumors exhibited a thicker fibrous capsule and a higher fibrous content with picnotic tumor cells and necrotic areas. Moreover, Ad*sflt*-treated tumors showed a less defined or non-existent peri-tumor hypervasculature border zone when compared to untreated or AdCMV.ßgal-injected tumors. When serial sections of mammary gland areas from Ad*sflt*-infected mice bearing regressed tumors were analyzed, areas of loose connective tissue containing stromal fibroblasts and some inflammatory cells were observed (Figure 3D). The presence of tumor cells remaining after treatment was detected by pan cytokeratin immunostaining; no positivity for tumor cell-specific anti-cytokeratin pan antibody in serial tissue sections from regressed tumor areas was observed (data not shown).

**Figure 3.**
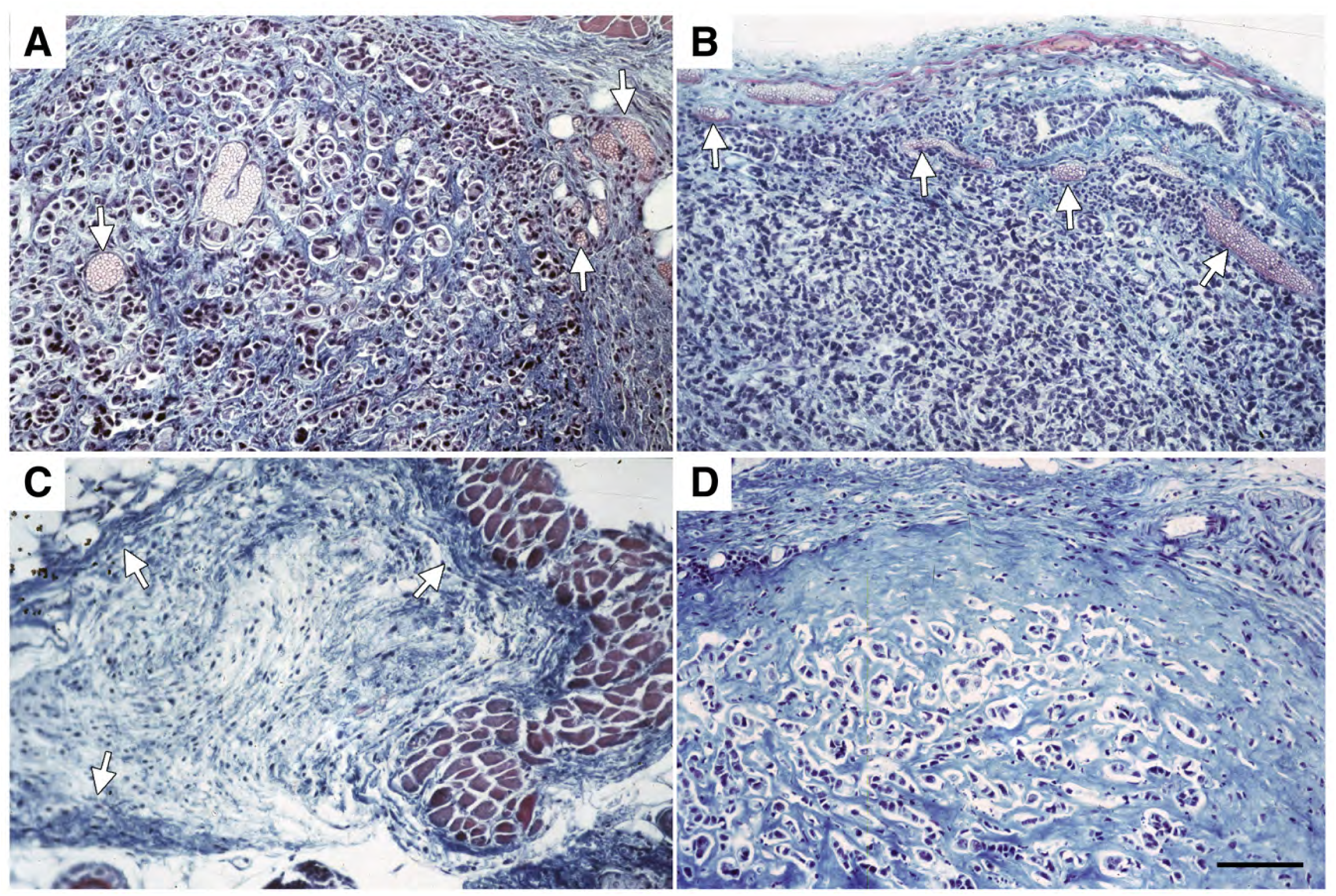
Histology of mammary tumors 6 months after Ad*sflt* injection. **(A)** Untreated tumors show intense vascularization (arrowheads) at d=0. **(B)** At 26 weeks AdCMVß gal-infected tumors consist of highly packed tumor cells with a thin fibrous capsule and a well-developed peri-tumor hypervasculature border zone (arrowheads). **(C)** Twenty six weeks after infection, Ad*sflt*-treated tumors are hypovascularised and exhibit a thicker fibrous capsule (arrowheads). The tumor is characterized by increased necrotic areas. **(D)** A tumor in regression after treatment with Ad*sflt* exhibits a high content of fibrous tissue and necrotic cells. All tumor sections were stained with Masson trichrome. The same magnification was used for all sections. Bar = 22⍰m.

### Expression of sFLT-1 in Adsflt-treated Tumors

To investigate whether the therapeutic effect on Ad-treated tumors *in vivo* was due to sFLT-1 expression, the extent and timing of Ad-transduced sflt-1 gene expression were analyzed by Northern blot and immunohistochemistry.

Remarkably high levels of sflt-1 mRNA expression were detected in Ad*sflt*-injected MCF-7 tumors from 3 days up to 6 weeks post-infection (Figure 4A). Transgene activity was still detectable, though sometimes at lower levels, 15 weeks after the Ad delivery. As expected, no endogenous sFLT-1 gene expression was detectable in AdCMV.βgal-treated tumors.

**Figure 4.**
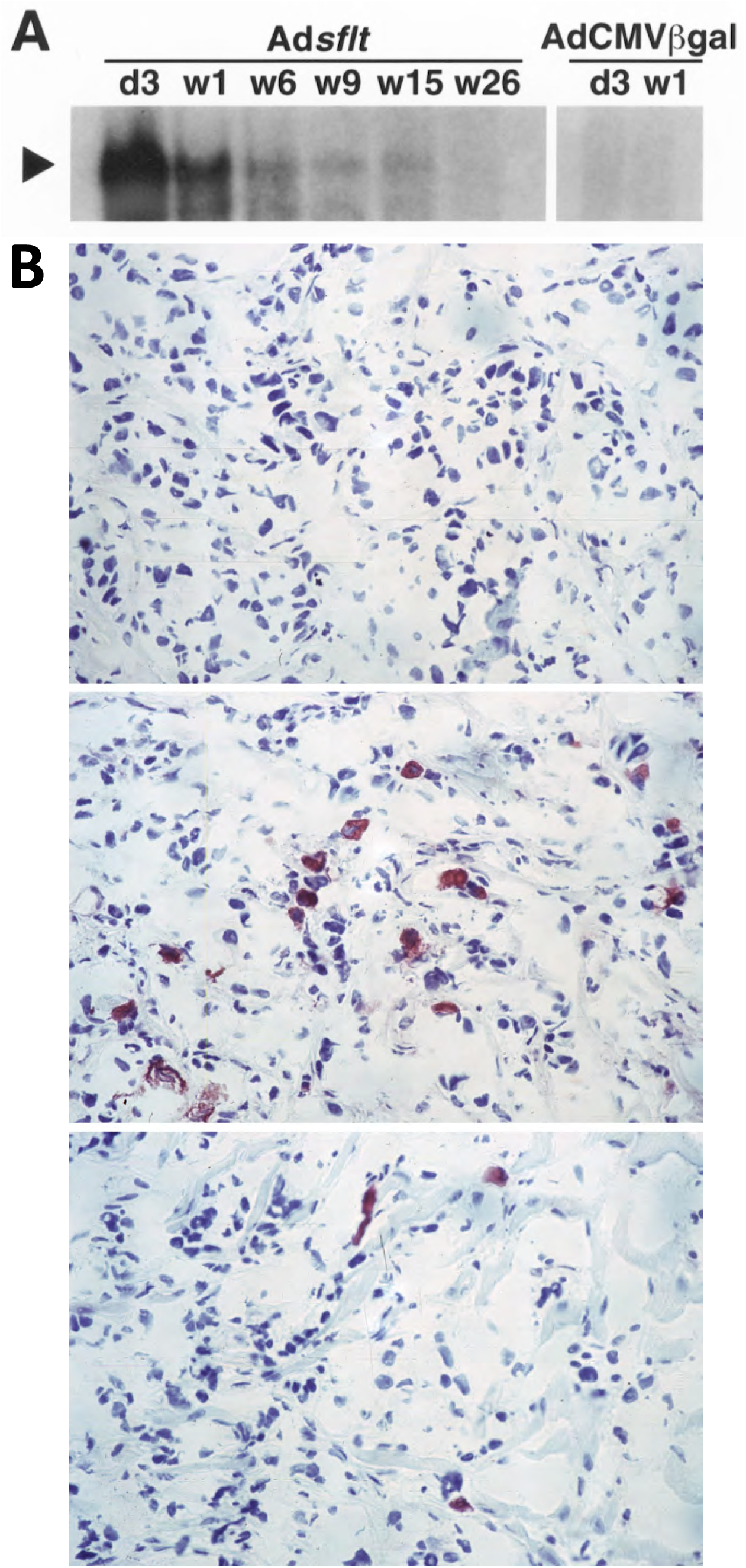
Analysis of sflt-1 mRNA and protein expression in Ad*sflt*-treated mammary tumors. **(A)** Northern blot analysis of sflt-1 mRNA expression *in vivo*. RNA was isolated from single Ad*sflt*-or AdCMV.βgal-treated tumors at the indicated time points. sflt-1 mRNA was markedly expressed in mammary tumors from day 3 up to 6 weeks after Ad*sflt* injection, and was still detectable 15 weeks after treatment. RNA from AdCMV.βgal-treated mammary tumors was used as negative control. *d*, day; *w*, week. **(B)** Immunohistochemical staining of representative sections from mammary tumors inoculated with control Ad virus (*top panel*) or Ad*sflt* (*middle and bottom panel*). Positive (red) staining indicates the expression of sFLT-1 protein in tumor sections, 7 days (*top and middle panel*) and 3 weeks (*bottom panel*) after treatment. The same magnification was used for all sections. Bar = 11 μm.

To confirm whether sFLT-1 protein was expressed during the development of infected tumors, we also performed immunohistochemical staining of paraformaldehyde-fixed, frozen tumor sections with an anti-sFLT-1 specific antibody. The results showed that while AdCMV.βgal-injected tumors exhibited no specific immuno-staining for sFLT-1 (Figure 4B), Ad*sflt*-injected tumors showed a specific sFLT-1 immuno-positivity localized to the cytoplasm of tumor cells, throughout the course of the experiment (Figure 4B). The most intense and homogeneous sFLT-1 staining was detected 7 days after treatment, and although the number of sFLT-1 immuno-positive cells gradually decreased, single positive cells were still found 26 weeks after Ad*sflt* injection (not shown).

### Adsflt Induces Vascular Regression in vivo and Inhibits Tumor Angiogenesis

Since the infection of MCF-7 cells with Ad*sflt* did not inhibit tumor cell growth *in vitro* (data not shown), Ad*sflt*-mediated inhibition of tumor growth was most likely due to the anti-VEGF activity of sFLT-1 *in vivo*, and not to a direct effect of Ad*sflt* on tumor cell growth rate. To determine whether the decrease in the volume of Ad*sflt-*treated tumors correlated with a decrease in vascular supply, tumor area and blood vessel area of similar-sized tumor sections for each time point were quantified on Masson trichrome-stained paraffin sections using an image analysis system. CD31 stained tumor sections were used to confirm the identity of small blood vessels. The blood vessel area to tumor area ratio (VA/TA) of Ad*sflt*-treated tumors (three sections from three tumors, n=9) and AdCMV.βgal-treated tumors (n=9) was similar 3 days post injection (0.385 ± 0.0702 *vs* 0.379 ± 0.0984; not significant) (Figure 5), but it was significantly reduced in the Ad*sflt* group from day 7 onward (*P* = 0.0199). Consistent with the inhibition of tumor angiogenesis, Ad*sflt*-treated tumors displayed 50% and 90% reductions in VA/TA at 3 weeks and 6 months, respectively, relative to control tumors (*P* <0.0001) (Figure 5). These results strongly suggest that sFLT-1 blocked tumor growth by inhibiting tumor angiogenesis.

**Figure 5.**
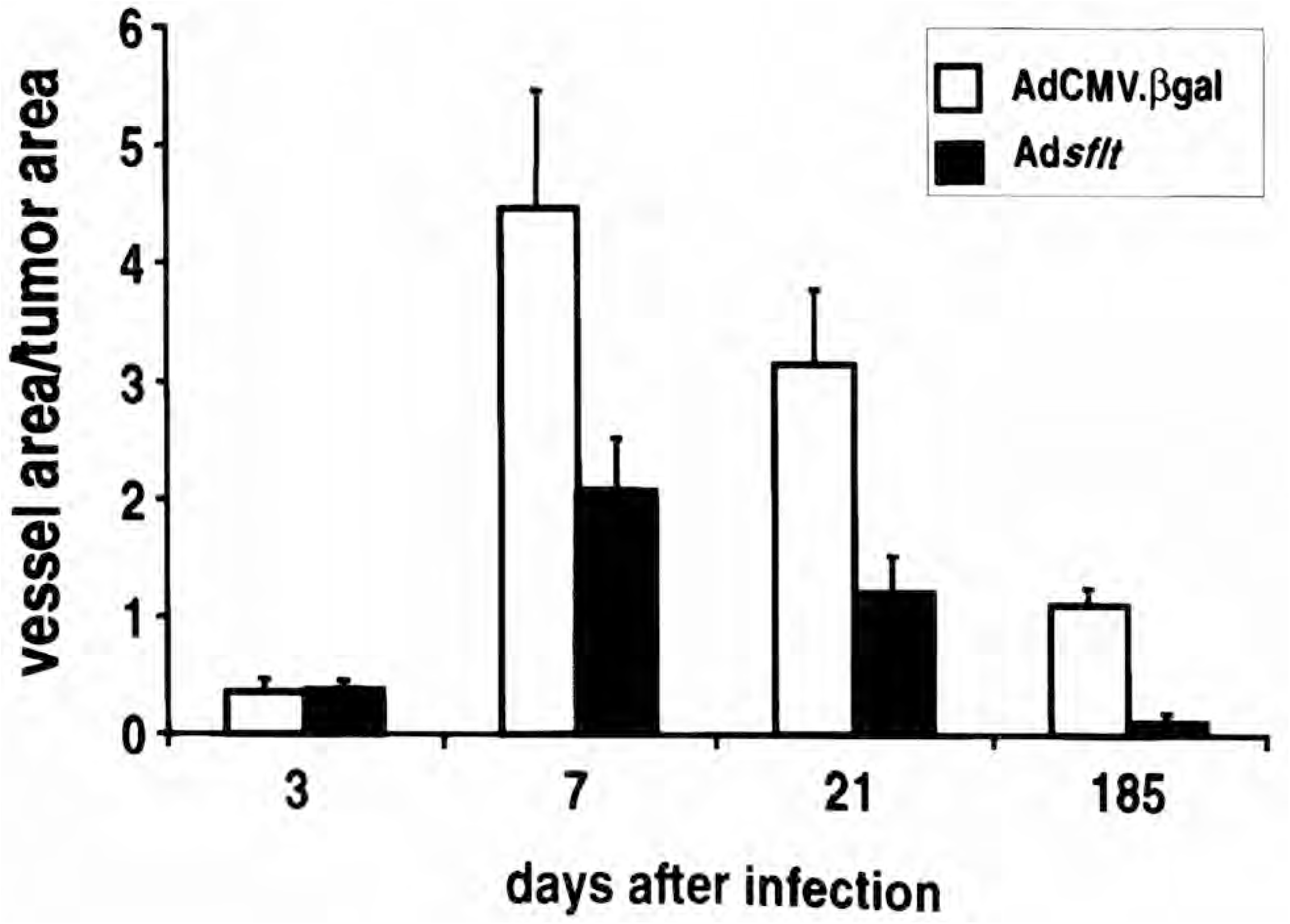
Ad*sflt* induces blood vessel regression *in vivo*. To determine whether Ad*sflt* could inhibit tumor vascularization, the blood vessel area/tumor area ratio was evaluated on Masson trichrome-stained tumor tissue sections using a KS300 imaging system. After 7 days, the tumors treated with Ad*sflt* showed a steady decrease in their vessel area/tumor area ratio. Six months after treatment (185 days), there was a 9.5 fold decrease in tumor vascular density in Ad*sflt*-compared to AdCMV.βgal-infected tumors. Graph shows mean values of vessel area/tumor area ± SE (n=9 per condition and time point) (*P* < 0.05).

### A Single Injection of Adsflt Induces Endothelial Cell Apoptosis

To further investigate the mechanism of sFLT-1 anti-angiogenic activity, a specific TUNEL test for apoptotic cell death was performed on Ad*sflt*-, AdCMV.βgal-or saline-treated tumor sections at various time points. The results showed a striking induction of apoptosis in vascular endothelial cells of Ad*sflt*-infected tumors as early as 7 days after injection (Figure 6). The same effect was still visible at 3 weeks after treatment (data not shown). Furthermore, positively stained endothelial cells were round and loosely attached to basement membrane. The study of serial sections from Ad*sflt* tumors confirmed that blood vessels harboring apoptotic cells showed signs of collapse and occlusion (Figure 6B and C). In contrast, AdCMV.βgal-or saline-treated tumors showed no apoptotic endothelial cells at any time point. CD31 stained tumor sections were used to confirm the identity of small blood vessels (data not shown).

**Figure 6.**
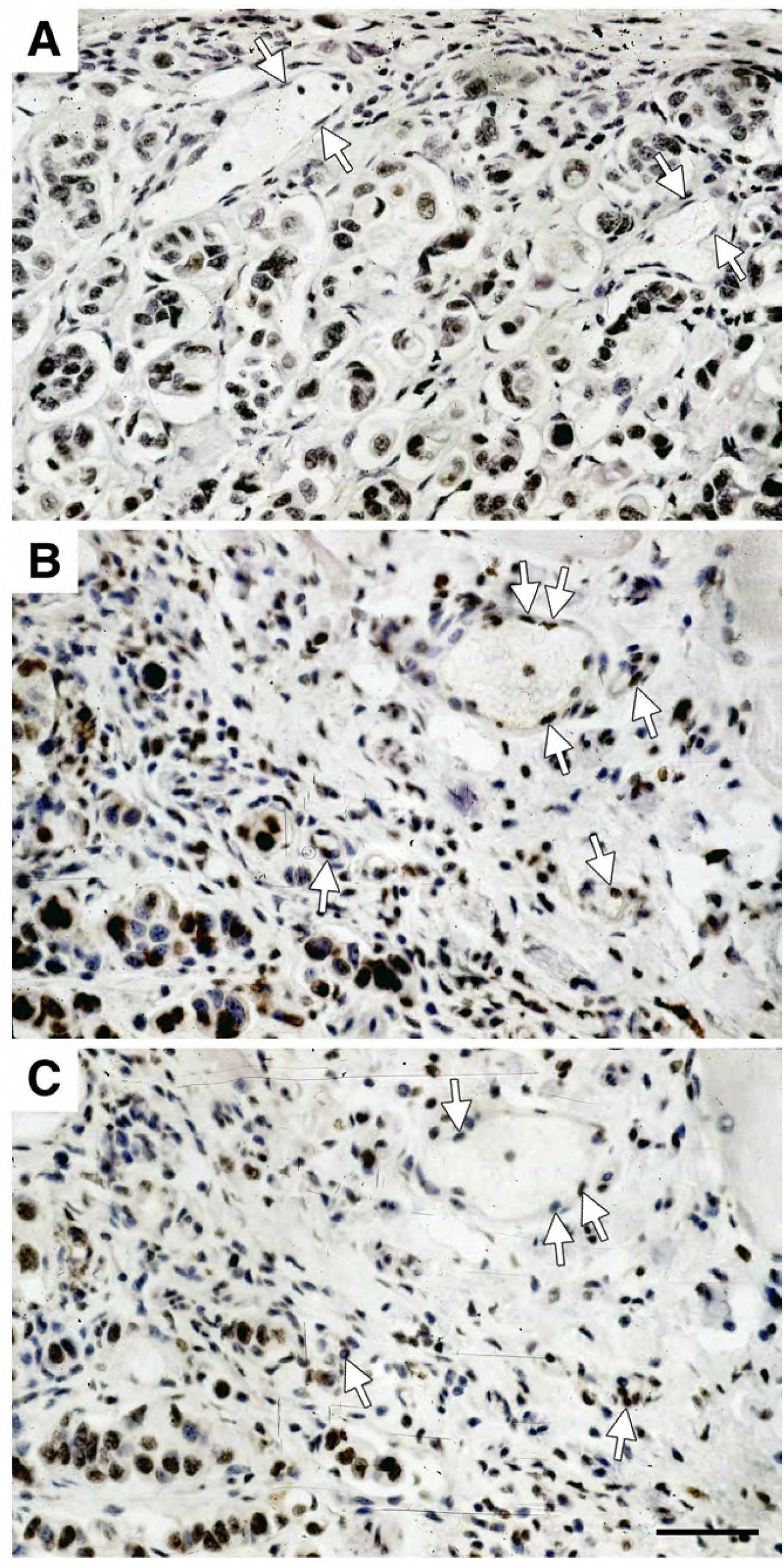
Ad*sflt*-mediated gene transfer induces tumor vascular endothelial cell apoptosis. Histologic sections from control and Ad*sflt* tumors were tested by TUNEL method to visualize apoptotic cells. Control Ad virus-injected tumors did not exhibit apoptotic reaction in endothelial cells (arrowheads) (A) at any time point; while apoptic reaction, indicated by brown nuclear staining, is visible in Ad*sflt*-injected tumor serial sections (B and C), 7 days after treatment. Note that apoptotic endothelial cells appear round and loosely attached to basement membrane (arrowheads). The same magnification was used for all sections. Bar, 11μm.

### Serum Levels of sFLT-1

The duration and magnitude of sFLT-1 expression in tumors, as well the impressive therapeutic effect observed, prompted us to determine whether serum levels of sFLT-1 were also increased in treated mice. Serum samples from untreated tumor-bearing mice and Ad*sflt*-or control Ad virus-infected mice (n=3 for each group) were analyzed by ELISA at different time points. As shown in Table 1, endogenous murine sFLT-1 could not be detected in untreated tumor-bearing nude mice, possibly due to poor recognition of murine sFLT-1 by the anti-human FLT-1 antibody 11G2, and/or to serum levels below the limit of ELISA detection. In contrast, human sFLT-1 was detectable only in Ad*sflt*-treated mice in the range of 0.2-6 ng/ml, with a marked increase 3 weeks after infection.

**Table 1.** Human sFLT-1 serum levels in nude mice bearing Ad vector-injected mammary tumors.

| Treatment | days<sup>a</sup> | sFLT-1 (ng/ml) |
| --- | --- | --- |
| untreated | 0 | nd<sup>b</sup> |
| Adcontrol<sup>c</sup> | 3 | nd |
| *Adsflt* | 3 | 2.47 ± 0.82 |
| Adcontrol | 21 | nd |
| *Adsflt* | 21 | 5.86 ± 5.62 |
| Adcontrol | 185 | nd |
| *Adsflt* | 185 | 0.23 ± 0.13 |

<sup>a</sup>days after intratumor injection of Ad vectors. <sup>b</sup>not detectable. The lowest detection limit of the assay is 0.2ng/ml. <sup>c</sup>Different transgenes encoding Ad vectors were used as negative controls with the same results. Data are presented as means + SE (n=3 mice per treatment group and time point).

### Regional Overexpression of sFLT-1 Does Not Inhibit Neovascularization During Wound Repair

The potential risk of systemic inhibition of angiogenesis is particularly important in anti-angiogenesis-based cancer gene therapy. To assess whether the observed increase of sFLT-1 serum levels in Ad*sflt*-treated animals could impair physiological angiogenesis, we studied wound healing following cutaneous injury 12 days after tumor infection (Brown *et al*., 1992; Frank *et al*., 1995). Full-thickness backskin excisions were made in 2-month-old female nude mice bearing Ad*sflt*-, AdCMV.βgal-or saline-injected tumors, and the decline in wound area over a two week period was measured. As shown in Figure 7, animals from all treatment groups showed comparable wound closure rates, with no observable delay of the Ad*sflt*-treatment group in the overall healing process. In all treatment groups most wounds reached 50% closure within 4 days, and all wounds closed between 10 and 14 days after injury. Thus, preliminary results suggest that a single injection of Ad*sflt* into tumors under the experimental conditions described above does not exert distant effects on wound healing.

**Figure 7.**
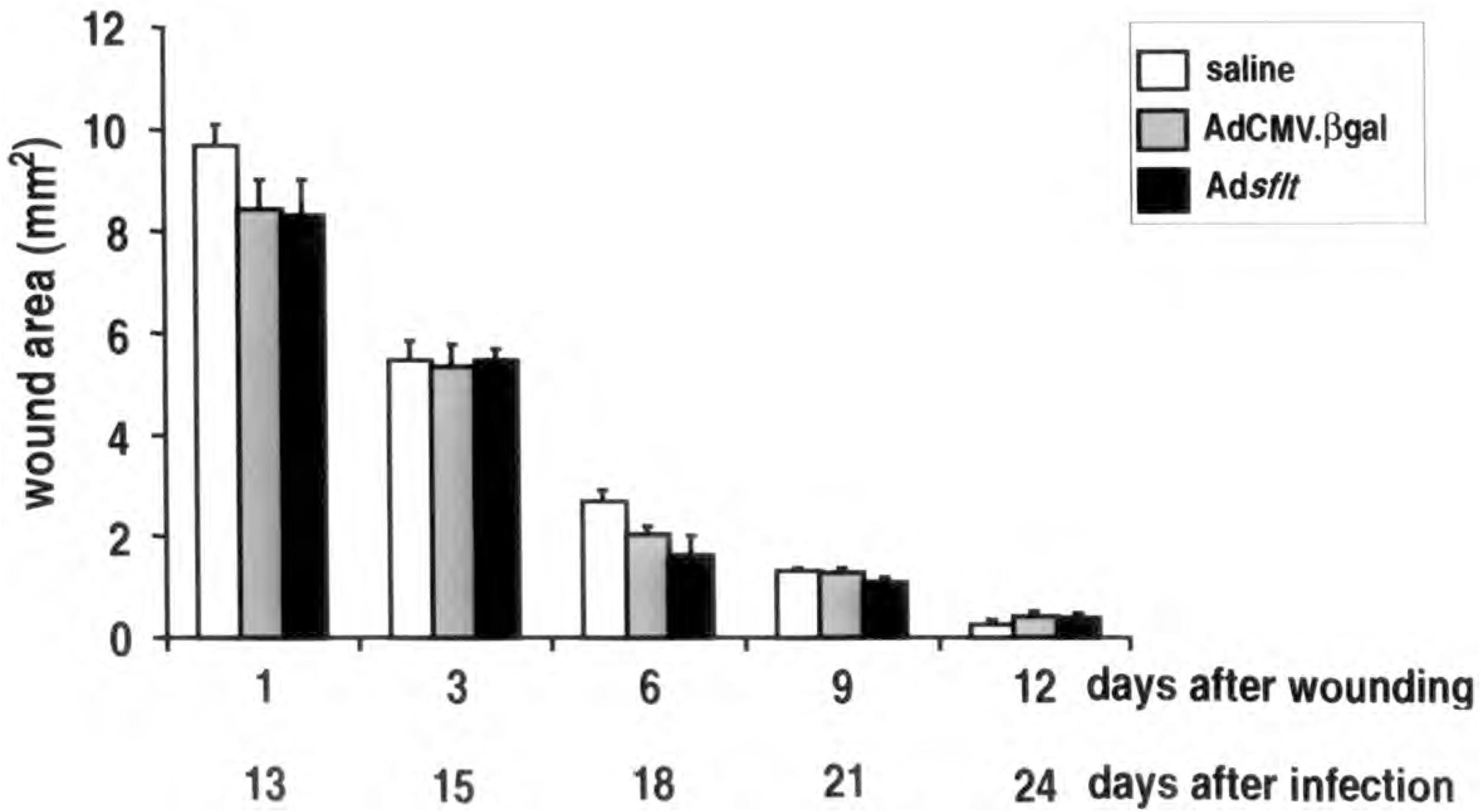
Ad*sflt* intratumor injection does not interfere with physiologic neoangiogenesis occurring during wound repair. Comparison of wound closure rates in nude mice injected intratumorally with saline, AdCMV.βgal or Ad*sflt*. One full-thickness skin excision was made on the upper dorsum of each female adult mouse and left undressed. At the indicated time points wound areas were measured. Graph shows mean wound area expressed as mm^2^ ± SE (n=6 mice per treatment group) (*P* < 0.05).

## Discussion

The present study demonstrates sustained anti-tumor effects of the Ad*sflt* vector in an orthotopic breast cancer model established in nude mice using the human MCF-7 carcinoma cell line. A single dose of Ad*sflt* vector resulted in 91% inhibition of tumor growth (i.e, a treated/control volume of 0.09) and a 67% regression of treated tumors over a period of six months. Following treatment, tumor growth inhibition and regression were quantifiable at the macroscopic level from week 7 onward. Northern blot and immunohistochemical analysis of Ad*sflt*-infected tumors revealed that the sflt-1 transgene was overexpressed by tumor cells, and that sFLT-1 protein was abundantly produced and secreted into the extracellular space. It is highly probable that VEGF present in this space would be tightly bound by the sFLT-1 protein, resulting in its sequestration from endogenous membrane-spanning VEGF receptors overexpressed on the tumor endothelium (Kendall *et al*., 1996) and VEGF signal blockade. sFLT-1 protein might also inhibit VEGF signaling by forming inactive heterodimers with endogenous VEGF receptors.

We also observed that wound healing, a physiological process which requires new blood vessel development, was not impaired in Ad*sflt*-treated tumor-bearing animals. This was of interest because anti-angiogenic strategies for the treatment of solid tumors pose the inherent risk of inhibiting physiological angiogenesis (Klauber *et al*., 1997). An important advantage of adenoviral vectors is their local activity. In the present study, the sflt-1 gene was transcribed in tumor cells, and the encoded protein secreted into the matrix present in the extracellular space. Elevated serum levels of sFLT-1 were detected in Ad*sflt*-treated tumor-bearing animals, but no impairment in the healing of full-thickness skin wounds, which requires physiological neovascularization, was observed. While a comprehensive investigation of this issue will require much additional experimentation, our results provide added support for the intratumoral expression of anti-angiogenic factors as a sound anti-cancer strategy.

Among anti-angiogenic therapeutic strategies, some divergence has been encountered regarding the duration of tumor growth inhibition, and our results are the first to show sustained inhibition using Ad*sflt*. Bazan-Peregrino et al. (2013) utilized AdEHE2F, which was designed to replicate in cells with high HIF and/or active estrogen signaling in the ER positive breast cancer cell line ZR75.1 They found that virotherapy using AdEHE2F-luc potently inhibited tumor growth *in vivo* but did not show that sFlt-1 could inhibit tumor angiogenesis. In our study, AdsFlt-1 substantially reduced ER positive breast cancer model MCF7 xenograft tumor growth in the presence of VEGF whereas our control (AdCMV.βgal and saline) treated mice failed to show inhibition of tumor growth.

Long-term tumor growth inhibition and regression has been demonstrated for malignant brain tumors with retroviral or adenoviral vectors encoding the thymidine kinase gene from herpes simples virus (Izquierdo *et al*., 1995; Maron *et al*., 1996), and with an adenoviral vector containing wild-type p53 in a squamous cell carcinoma (Pirollo *et al*., 1997) and triple-negative breast cancer (Wang et al, 2015). Using endostatin, complete tumor regression could be achieved after several treatment cycles (Boehm *et al*., 1997). A prior study using a plasmid expression vector encoding sFLT-1 in an *ex vivo* tumor model showed that tumor growth was inhibited over a period of 2-3 weeks (Goldman *et al*., 1998). Another independent study reported inhibition of tumor growth following the regional or intratumoral expression of Ad*sflt* over 2 weeks (Kong *et al*., 1998). The striking duration of tumor regression and the lack of recurrence over a six month period we report may be as much attributable to the modestly aggressive MCF-7 cell tumor model as it is to the efficacy of the Ad*sflt* vector or the mechanism of action of sFLT-1 protein. Nonetheless, the apoptotic effect of sFLT-1 on endothelial cells overcomes the main disadvantage of adenoviral vectors, namely their transient expression. The very rapid anti-angiogenic effect seen as early as 7 days after treatment suggests that under the conditions used here, sustained high-level expression of sFLT-1 was not critical to its overall efficacy.

We demonstrate that the ability of sFLT-1 to induce tumor regression correlated with a significant decline in the number of blood vessels. In fact, within 7 days of infection, Ad*sflt*-treated tumors displayed three important features characteristic of anti-angiogenic therapy: i) diminution of VA/TA ratio; ii) apoptosis of endothelial cells; and iii) necrosis of tumor cells. Since the quantification of blood vessel area per tumor area showed that blood vessel regression was independent of tumor size, we conclude that the general basis of Ad*sflt* activity *in vivo* was a paracrine effect on tumor angiogenesis. While the precise mechanism by which inhibition of VEGF signaling results in this sequence of events remains to be elucidated, several highly relevant observations have been reported. In addition to its mitogenic effect on endothelial cells, VEGF is also known as a survival factor for endothelial cells (Alon *et al*., 1995; Melder *et al*., 1996). A critical level of VEGF activity for endothelial cell survival has been demonstrated by the heterozygous lethality of the VEGF knockout mice (Carmeliet *et al*., 1996; Ferrara *et al*., 1996), and in androgen-dependent tumors, where decreased VEGF expression after androgen withdrawal leads to vessel regression and apoptosis of tumor endothelial cells (Jain *et al*., 1998). In addition, induction of apoptosis in endothelial cells in the absence of growth factors has been shown *in vitro* by angiostatin, an endogenous inhibitor of neoangiogenesis (O’Reilly *et al*., 1994), and in xenografted C-6 glioma cells by a tetracyclin-regulated VEGF expression system. In this animal model, endothelial cells detach from tumor blood vessels and undergo subsequent death by apoptosis when VEGF production is turned off (Benjamin and Keshet, 1997). Thus it is likely that in our model, VEGF binding to sFLT-1 results in locally decreased VEGF signaling, leading in turn to endothelial cell apoptosis.

Consequent to sFLT-1-mediated inhibition of angiogenesis and endothelial cell apoptosis, striking tumor cell necrosis resulted in tumor regression in almost 67% of the Ad*sflt* -treated tumors. Serial tissue sectioning and immunohistochemistry revealed that in animals where we could find residual tumor mass, it consisted almost entirely of fibrous tissue, whereas control animals showed typical breast carcinomas. Again, although the precise mechanism by which this occurs is not yet known, VEGF enhances vascular permeability generally (Dvorak *et al*., 1995), and renders tumor vessels hyperpermeable to plasma proteins and other macromolecules, allowing their escape into the tumor matrix. VEGF may provide a matrix enriched with nutrients and growth factors which facilitates tumor growth.

Consistent with this notion, tumor vascular permeability can be reduced by neutralization of endogenous VEGF (Yuan *et al*., 1996). sFLT-1-mediated blockade of VEGF signaling in our model may have led to tumor cell necrosis through reduced vascular permeability and consequent reduction in the supply of needed nutrients and growth factors. Interestingly, the residual tumor dormancy characteristic of other anti-angiogenic factors as angiostatin (O’Reilly *et al*., 1994) was not observed in our study of sFLT-1-mediated tumor regression.

In summary, anti-angiogenic therapy using a recombinant adenovirus encoding sFLT-1 was found to be highly effective in the MCF-7 breast carcinoma animal model. Ad*sflt* may be well-suited for the treatment of solid tumors with moderate growth rates because of the rapid and strong anti-angiogenic action of sFLT-1. The complete tumor regression in the absence of obvious inhibition of physiological angiogenesis demonstrated here suggests that this promising anti-cancer strategy deserves considerable further investigation.

## Materials and Methods

### Cell Culture

Human umbilical vein endothelial cells (HUVEC, Clonetics, San Francisco, CA) were maintained in complete endothelial cell growth medium (EGM-2, Bio-Whittaker, Walkersville, MD) containing endothelial cell basal medium (EBM-2, Bio-Whittaker) supplemented with endothelial cell Bullet Kit (Bio-Whittaker). Cells were used for experiments between passages 3 and 5. MCF-7 human breast carcinoma cells (ATCC, Rockville, MD) were grown in Dulbecco’s MEM (DMEM) with Glutamax-1 (Gibco, Paisley, Scotland) supplemented with 10% fetal bovine serum (FBS), penicillin (100 U/ml) and streptomycin (100 µg/ml), and used for injection into nude mice between passages 10 and 15.

### Adenoviral Vectors

For *in vitro* and *in vivo* studies recombinant, replication-deficient first generation adenoviral (Ad) vectors were used (Rosenfeld *et al*., 1991; Rosenfeld *et al*., 1992). Ad*sflt* expressed a 688 amino acid product comprising the secretory leader sequence and the amino-terminal six extracellular Ig-like domains of the full length human FLT-1 receptor plus a 31 amino acid carboxy-terminal tail specific to the secreted soluble form of the receptor (Kong *et al*., 1998). AdCMV.ßgal was used as a control vector (Maeda *et al*., 1994). In both vectors transgene transcription was driven by the cytomegalovirus (CMV) early promoter/enhancer. Ad vectors were amplified in 293 competent cells, purified by three centrifugations on a cesium chloride gradient, dialyzed, and stored at -80° C until use. Viral titers were determined by plaque assay using 293 cells. Viral stocks that became contaminated with replication competent wild type virus were excluded by testing for plaque forming capacity on A549 cells (Lochmuller *et al*., 1994).

### Proliferation Assay

HUVEC were plated in 35 mm petri dishes at a density of 25,000 cells/dish in EGM-2 medium. After 12 h, cells were counted (time point 0) and infected either with 25 plaque forming units (pfu)/cell of AdCMV.ßgal, Ad*sflt*, or were left uninfected. The infection was carried out in EBM-2 medium plus gentamycin, heparin and ascorbic acid (from the Bullet kit) for 90 min at 37° C with occasional rocking. After infection cells were cultured with EBM-2 containing 10% FBS, gentamycin, heparin and ascorbic acid in the presence or absence of human recombinant VEGF (10 ng/ml; R&D, Minneapolis, MN) or bFGF (2 ng/ml; R&D) as specified in Results. The culture medium was changed on days 2 and 4 after infection. Cells were counted with a hemocytometer in triplicate on days 1, 3, 4 and 5. The experiment was repeated twice.

### Animal Studies

All animal studies described were carried out in accordance with the guidelines set forth in the <u>NIH Guide for the Care and Use of Laboratory Animals</u> (1985). For the orthotopic breast cancer model, two-week-old female nude mice (CD1 nu/nu, Harlan-Italy, Correzzana, Italy) were inoculated with MCF-7 cells (1×10^6^/50 l of DMEM) into the fat pad of the mammary gland. Injections were carried out on both sides of the hind leg in the mammary gland area. After 5-6 weeks, when palpable solid tumors had developed, animals were randomly assigned to treatment groups, and both tumors of each mouse were injected with Ad vectors or saline solution. A single dose (2×10^9^ pfu/50 µl) of Ad*sflt* or AdCMV.ßgal or 50 µl of saline solution was injected locally into and around the tumor. Tumor size was measured with a caliper prior to injection and at weekly intervals thereafter. Tumor volume was calculated using the following formula: tumor volume (mm^3^) = a x b x c; where a, b, and c are the three measurable axes of the tumor. Tumors where excised for histological analysis or RNA extraction at days 0 and 3, and weeks 1, 3, 6, 9, 15 and 26. Since several tumors in the control groups had ulcerated the overlying derma, the experiment was terminated by sacrificing the animals 26 weeks after treatment. The animal study was repeated once following the same protocol and in both experiments tumor development showed a similar growth curve.

For wound healing assays, twelve days after intratumor injection with Ad vectors or saline solution, female nude mice (2-month-old CD1 *nu*/*nu* from Harlan Nossan) were anesthetized, their mid-dorsum were cleansed and a single full-thickness 4-mm diameter excisional wound (12.5 mm2) was made with a biopsy punch. The wound was left undressed and its area recorded on days 1, 3, 6, 9 and 12. The experiment was repeated twice (n = 6 animals per treatment group).

### Immunohistochemistry

sFLT-1 protein expression was evaluated on 3 µm frozen tissue sections, fixed in 4% paraformaldehyde, from control and Ad*sflt*-treated tumors using the rat anti-human sFLT-1 antibody 7A6 (Barleon *et al*., 1997). Sections were washed in PBS and pre-incubated in normal rabbit serum followed by incubation with 7A6 antibody (5 g/ml) at room temperature for 2 h and washed with PBS. A biotinylated anti-rat antibody (Vector Laboratories, Burlingame, CA) was used as secondary antibody at a dilution of 1:500 for 1h. Endogenous peroxidases were blocked by incubating the sections in 1% H_2_O_2_ for 1 h before applying a streptavidin-peroxidase complex (Vector Laboratories). The enzyme reaction was developed with 3-Amino-9-ethyl-carbazole and reinforced by incubating the sections with 0.5% copper sulfate solution. Sections were counter-stained with hematoxylin and mounted in aqueous medium.

Apoptosis was studied using an *in situ* cell death detection kit (Boehringer-Mannheim, Monza, Italy) on routine formaldehyde fixed paraffin sections. Terminal deoxynucleotidyl transferase-labeling was performed according to manufacturer’s instructions with the exception that sections were not pretreated with proteinase K. Sections were analyzed and photographed with an Axioplan 2 microscope (Zeiss, Jena, Germany).

Endothelial cells in tumor blood vessels were identified immunologically on 3 -5 µm thick formaldehyde-fixed paraffin sections that were pretreated twice in a microwave oven in 10 mM citrate buffer at 650 W. Sections were allowed to cool down to room temperature, and were then transferred to PBS and quenched with a peroxo-block solution (Zymed Laboratories, San Francisco, CA) for 5 min. After washing with PBS, sections were incubated with anti-PECAM-1 (CD31) antibody (Santa Cruz Biotechnology, Santa Cruz, CA) at a dilution of 1:200 at room temperature for 1 h. Incubation with a donkey anti-goat secondary antibody HRP conjugate (Santa Cruz Biotechnology) and 3-Amino-9-ethyl-carbazole was used to develop the enzymatic reaction. Sections were counter-stained with hematoxylin and mounted in aqueous medium. Anti-cytokeratin pan antibody (Boehringer-Mannheim) was used according to the supplier’s instructions.

### Northern Blot Analysis

For the detection of sflt-1 mRNA expression, total RNA was isolated from HUVEC or excised mammary tumors using the TRIZOL Reagent (Gibco BRL, Paisley, Scotland). The RNA concentration was determined spectrophotometrically and normalized by ethidium bromide staining of agarose gels. Twenty five g of total RNA was separated on 1% agarose-formaldehyde gels, transferred onto nylon membranes (Sartorius, Göttingen, Germany), and hybridized with 32P-labeled 450-bp sflt-1 cDNA probe. The sflt-1 probe was generated by polymerase chain reaction using the 5’ flt-1 (nt 1808-1827; 5’-GCACCTTGGTTGTGGCTGAC-3’) (Kendall and Thomas, 1993) and 3’ sflt-5 (nt 2238-2258; 5’-GAGATCCGAGAGAAAACAGCC-3’) primers and Ad*sflt* DNA as template. Membranes were hybridized with 1×106cpm/ml of QuikHyb solution (Stratagene, La Jolla, CA), washed once with 2xSSC/0.1% SDS for 15 min at 23°C, then twice with 2xSSC/0.1% SDS for 30 min at 63°C, and exposed to Kodak Bio-Max film (Eastman Kodak, Rochester, NY) at -80°C with intensifying screens overnight (HUVEC RNA analysis) or for 2 days (tumor RNA analysis).

### Quantification of Tumor and Blood Vessel Areas

To determine tumor and blood vessel areas, Masson trichrome-stained tissue sections were visualized with an Axioplan 2 microscope (Zeiss) using a 5x or 40x objective respectively, and total tumor area per tissue section, and total blood vessel area per tissue section, were measured with a KS300 imaging system (Kontron, Elekronik, Munich, Germany). Each tumor was cut in serial sections and 3 random central sections were analyzed. At least 3 tumors for each treatment group and time point were analyzed (n=9). For statistical evaluation only similar sized tumor areas of differently treated groups were analyzed at specific time points.

### sFLT-1-specific ELISA

Blood samples were obtained by abdominal artery puncture from anaesthetized mice before sacrifice, incubated at 37°C for 30 min and spun at 2,000 rpm at room temperature for 5 min. Serum samples were frozen at -80° C and thawed once before use. Serum sFLT-1 measurement was performed by ELISA as previously described (Barleon *et al*., 1997a) with minor modifications. Mouse monoclonal antibody 11G2 as capture antibody was used at 2µg/ml. For detection, a polyclonal rabbit anti-sFLT-1 serum was used at a 1:1500 dilution. For calibration purified recombinant sFLT-1 was used.

### Statistical Evaluation

Results are reported as mean ± standard error of the mean (SE). Statistical analysis was performed by two way analysis of variance using the Student-Newman-Keuls Test for pairwise multiple comparison of cell numbers, tumor volumes and wound areas. A one way analysis of variance was used comparing tumor area/blood vessel area ratio of similar sized tumor sections and a Student’s *t*-test was used for sFLT-1 serum level quantitation. Differences were considered statistically significant at *P* < 0.05.

## Acknowledgments

We thank Cesare M. Secci and Augusto Mari for their technical support and Gabriella Ricci for her excellent secretarial assistance during the preparation of this manuscript.

## Abbreviations

Ad: adenovirus
CMV: cytomegalovirus;
sFLT-1: soluble FLT-1;
VEGF: vascular endothelial growth factor

